# BRAF^V600E^ expression in thyrocytes causes recruitment of immunosuppressive STABILIN-1 macrophages

**DOI:** 10.1101/2022.07.27.501733

**Authors:** C Spourquet, O Delcorte, P Lemoine, N Dauguet, A Loriot, Y Achouri, M Hollmén, S Jalkanen, F Huaux, S Lucas, P Van Meerbeeck, JA Knauf, JA Fagin, C Dessy, M Mourad, P Henriet, D Tyteca, E Marbaix, CE Pierreux

## Abstract

Papillary thyroid carcinoma (PTC) is the most frequent histological subtype of thyroid cancers (TC), and BRAF^V600E^ genetic alteration is found in 60% of this endocrine cancer. This oncogene is associated with poor prognosis, resistance to radioiodine therapy and tumor progression. Histological follow-up by anatomo-pathologists reveals that 2/3 of surgically-removed thyroids do not present malignant lesions. Continued fundamental research into the molecular mechanisms of TC downstream of BRAF^V600E^ remains thus central to better understand the clinical behavior of these tumors.

To study PTC, we used a mouse model in which expression of BRAF^V600E^ is specifically switched on in thyrocytes by doxycycline administration. Upon daily intraperitoneal doxycycline injection, thyroid tissue rapidly acquired histological features mimicking human PTC. Transcriptomic analysis revealed major changes in immune signaling pathways upon BRAF^V600E^ induction. Multiplex immunofluorescence confirmed the abundant recruitment of macrophages, among which a population of LYVE-1+/CD206+/STABILIN-1+ was dramatically increased. By genetically inactivating the gene coding for the scavenger receptor STABILIN-1, we showed an increase of CD8+ T cells in this *in situ* BRAF^V600E^ dependent TC. Finally, we demonstrated the presence of CD206+/STABILIN-1+ macrophages in human thyroid pathologies. Altogether, we revealed the recruitment of immunosuppressive STABILIN-1 macrophages a PTC mouse model and the relevance of these observations in human thyroid tissues.

## INTRODUCTION

In recent years, thyroid cancer (TC) has attracted more and more attention due to the rapid increase in its incidence. TC ranks 5^th^ in incidence among all malignancies and is the most common endocrine tumor. In addition, the incidence rate is almost three times higher in women than in men [1]. The number of TC cases continues to increase not only due to external factors but mostly because of overdiagnosis of TC due to the higher sensitivity of diagnostic techniques [2]. Papillary thyroid carcinoma (PTC) accounts for 80-85% of TC cases [3] and about half of them carry the BRAF^V600E^ mutation [4,5]. The presence of this mutation is often associated with poor prognosis and distant metastasis. Although the 5-year survival rate is close to 98% when the cancer is diagnosed at an early stage, the survival rate drops to around 50% among people diagnosed with poorly differentiated thyroid cancer (PDTC) and/or distant-stage disease [6]. Another important challenge is the overdiagnosis of nodules or indolent microcarcinomas which leads to overtreatment and unnecessary surgeries [7,8]. It is thus important to better understand and characterize the biology and the microenvironment of TC to improve the diagnosis and treatment especially when the cancer is poorly differentiated.

Solid tumors like TC are composed of various populations of cells including tumor cells, cancer stem cells, stromal cells and immune cells. These cells together with the extracellular matrix and the molecules produced constitute the tumor microenvironment (TME), which plays a central role in tumor development and progression [9]. Indeed, in the TME, immune cells are supposed to recognize and eliminate cancer cells, but they rather progressively modify the TME to facilitate tumor progression. Tumor-associated macrophages (TAMs) are key components of the immune TME and are highly plastic cells with multiple functions [10]. TAMs are derived from tumor-infiltrating monocytes in peripheral blood or from expansion of tissue-resident macrophages [11]. TAM population can transition between pro-inflammatory and anti-inflammatory states [12], and this polarization status depends on the pathological context and environment. Pro-inflammatory macrophages display antitumor effects such as the identification of tumor cells and their elimination. Anti-inflammatory/immunosuppressive macrophages are involved in tissue fibrosis and remodeling, angiogenesis, tumor progression, and immunoregulation towards immunosuppression [13].

The composition of the TC microenvironment depends on the mutations carried by the tumor cells, the cell type initiating the tumor, and the differentiation status of the tumor [14]. By considering the immune infiltration pattern of the TME and the gene expression profiling, a new classification of TC has been proposed. Two different clusters, the anaplastic thyroid cancer (ATC) and the PDTC, have been described among which PTC display a mixed immunological behavior with these two clusters [15]. Another study reveals that the presence of CD4+ T cells correlates with tumor size and aggressive features of TC while the presence of cytotoxic CD8+ T cells in the TME is associated with a favorable prognosis in patients with PTC [16]. A reduced CD4+/CD8+ T cell ratio is thus indicative of a good prognosis. TAM density also positively correlates with larger tumor size, lymph node metastasis and decreased survival in PTC [17,18]. In a mouse model, PTC tumor expressing BRAF^V600E^ show high TAM infiltration in response to increased expression of TAM chemoattractant, colony stimulating factor-1 (CSF-1) and chemokine CCL2, by cancer cells [19]. In this mouse model and in TC in general, TAMs display an immunosuppressive phenotype [19,20]. Despite these important pieces of information on TC microenvironment, the complete identity of the immune microenvironment as well as the complex and dynamic crosstalk between TC cells and their TME still needs to be unraveled.

In this study, we aimed to better understand TME cellular composition, evolution and role using a Tg-rtTA and tetO-BRAF^V600E^ double transgenic mouse model, mimicking BRAF^V600E^-dependent PTC. Total RNA sequencing of thyroid tissue revealed major changes in different immune signaling pathways, as soon as two days after BRAF^V600E^ induction. Among the most deregulated genes, we found several chemokines and their receptors, which are mainly expressed by macrophages. By immunohistofluorescence staining, we characterized these macrophages and identified a particular population which expressed LYVE-1 (a lymphatic vessel-specific glycoprotein), CD206 and STABILIN-1 (a scavenger receptor) markers. Using CRISPR-Cas9, we inactivated *Stabilin-1*, and found that its absence did not affect tumor size nor epithelial characteristics but was associated with a decrease in the intratumoral CD4+/CD8+ T cell ratio, thereby supporting a potential immunosuppressive role for the STABILIN-1 macrophages. Finally, we extended our observations to human tissues and showed the presence of these macrophages on sections from patients with benign and malignant thyroid pathologies.

## MATERIALS & METHODS

### 1. Animals

Tg-rtTA and tetO-BRAF^V600E^ mice were obtained from J. A. Fagin [21]. Mice were intraperitoneally injected with a saline solution (CTL) or with doxycycline (1µg/g body weight) every 24h during maximum 4 days (Day 1 → Day 4) [22]. The *Stabilin-1* knockout mice were generated at the UCLouvain Transgenesis platform as described [23]. Sequences targeted by the crRNAs (crRNA1: 5’-AGGAAAGAATTCAGAACTGG-3’; crRNA2: 5’-GTGATCGTTACCTATCCTCC-3’) were chosen to enclose the transcriptional initiation site and the entire first exon (Figure 3A). Pups were screened for the deletion by classical genotyping PCR with the GoTaq R2 Hot Start Green Master Mix (Promega) with fw (5’-CCTCGGAAGCTGCCTAAGAT-3’), rv (5’-AAGGGAAAATGTACGGACACG-3’), fw’ (5’-TGTGGGGACAGTGATTGCAG-3’) and rv’ (5’-CCACCCACCACAAGCATAGA-3’) primers in the presence of DMSO 5% (Figure 3B). Three heterozygous males (FVB mix B6D2) and four heterozygous females (FVB mix B6D2) were selected based on Sanger sequencing of the PCR products and of a ± 1300bp deletion to initiate the Tg-rtTA; tetO-BRAF^V600E^; *Stabilin-1* knockout colony. This colony was further backcrossed into FVB background. Mice were raised and treated according to the NIH Guide for Care and Use of Laboratory Animals, and experiments were approved by the University Animal Welfare Committee of the UCLouvain, Brussels, Belgium (2016/UCL/MD/005 and 2020/UCL/MD/011).

### 2. Tissue collection

Mice were anaesthetized by injection with a xylazine (20mg/kg)/ ketamine (200mg/kg) solution and sacrificed by cervical dislocation after blood puncture via the retro-orbital vein and cardiac flushing with PBS. Thyroid lobes were excised and either snap-frozen for RNA analysis or cut in 1mm^3^ pieces for tissue dissociation and flow cytometry analysis. For histological or immunolabelling analysis on paraffin sections, thyroid lobes were extracted with the trachea, fixed in 4% paraformaldehyde at 4°C overnight and embedded in paraffin using a Tissue-Tek VIP-6 (Sakura). Sections of 6µm were obtained with the microtome Micron HM355S (ThermoScientific). For immunohistofluorescence on frozen sections, thyroid lobes were collected, fixed in 4% paraformaldehyde at 4°C overnight, followed by an overnight immersion in PBS/20% sucrose solution. Finally, tissues were embedded in PBS/15% sucrose/7.5% gelatin solution and stored at -80°C. Sections of 7µm were obtained with a cryostat (Thermo Scientific, Cryostar NX70). Frozen human tissue samples (sections or tissue pieces) were obtained from the Saint-Luc Hospital’s biobank, Brussels. The study followed the Declaration of Helsinki and was approved by the local ethics committee of the UCLouvain, Brussels, Belgium (2017/25OCT/495 and 2017/04OCT/466).

### 3. Histology and immunohistochemistry

After paraffin removal and rehydration, sections were either stained with hematoxylin and eosin for histological analysis or treated with 0.3% H2O2 for immunohistochemistry. Slides were processed as described [22] for antigen retrieval, permeabilization, blocking and antibody staining (listed in Supplementary Table S1). For both, histology and immunohistochemistry, slides were mounted in Dako aqueous Medium (Agilent Technologies) and scanned with the panoramic P250 digital slide scanner (3DHistech).

### 4. Immunohistofluorescence and image analysis

For LYVE-1, CD11b, CD206, STABILIN-1 labelling, frozen sections were used. Tissues were treated as described [23] for permeabilization, blocking and staining steps. For F4/80, LAMP-1 and TG staining, paraffin sections were used. After paraffin removal and rehydration, sections were treated as for immunohistochemistry (see above) for antigen retrieval, permeabilization, blocking and labelling with primary antibodies (listed in Supplementary Table 1). For CD3 and CD8 labelling, tissues on paraffin sections were treated as described [24]. Sections were then blocked with PBS/0.3% triton X-100/10% BSA/3% milk solution for 45min, and incubated with anti-E-CADHERIN antibody overnight at 4°C. Secondary antibodies coupled to Alexa-488, -568 or -647 (Invitrogen) were used as well as a fluorescent nuclear dye (Sigma, Hoechst 33258). Slides were finally mounted with Dako Fluorescent Mounting Medium, and scanned with the Pannoramic P250 Digital Slide Scanner (3DHistech), or observed with the Zeiss Cell Observer Spinning Disk (COSD) confocal microscope. Images were analysed with Zen software (Zeiss). Image quantifications of CD3 and CD8 cells were performed with the HALO software and its Cytonuclear FL module (Indicalabs). Due to the lack of a validated anti-CD4 antibody on tissue sections, CD4+ T cells were considered as CD3+ CD8-cells and CD8+ T cells as CD3+ CD8+ cells.

### 5. Blood analysis and ELISA

Blood was collected with sodium-heparinized capillary tubes from the retro-orbital sinus. Blood was analysed by the MS9-5s hematology analyzer (MS laboratoires). For ELISA, blood was centrifuged at 1880g for 10min to collect the plasma fraction. Plasma was then centrifuged at 2500g during 10min to eliminate remaining red blood cells and platelets. Plasma TNFα was quantified using mouse TNFα uncoated ELISA kit (Invitrogen) according to the manufacturer’s instructions.

### 6. mRNA quantification

Total RNA was extracted from mice thyroid tissues using TRIzol Reagent (Thermo Scientific), as described in [25] followed by an additional phenol/chloroform purification step. Total RNA was extracted from human thyroid tissues using miRNeasy Mini kit (Qiagen) according to the manufacturer’s instructions and as described [26]. Tissue-RNA concentration was measured by spectrophotometry (Thermo Scientific, NanoDrop 8000). 500ng of total RNA from each sample were reverse-transcribed using M-MLV Reverse Transcriptase (Invitrogen) and random hexamers. Real-time quantitative PCR (RT-qPCR) was performed as described [27]. Primer sequences are listed in Supplementary Table S2. Data were analysed using the ΔΔCT method, with the geometric mean of *Gapdh* and *Rpl27* expression as reference for mice tissues, and *RPL13A* expression as reference for human tissues.

### 7. mRNA sequencing and bioinformatic analysis

DNase-treated RNA samples from control (CTL), Dox 2 days (Day 2) and Dox 4 days (Day 4) thyroid (n=4; 2 males and 2 females for each group) were sequenced by GENEWIZ-NGS Europe (Germany) using Illumina NovaSeq system. Raw data in fastq format were processed with R software. According to the p-value histogram and the PCA plot, all samples were used for differential expression analysis. Enrichment pathway analyses were performed with MouseMine software.

### 8. Flow cytometry

Spleen and thymus were crushed with the back of a 5ml syringe in 5ml of IMDM medium (Gibco, 21980-032) supplemented with 50µM β-mercaptoethanol (Gibco, 31350-010), glutamax and 5% FBS. Cell suspension was filtrated on a 70µm-filter and centrifuged at 300g for 6min. Splenic and thymic cell suspension, as well as blood samples, were resuspended in 4ml of Red Blood Cell lysis buffer (eBioscience, 00-4300-54) for 5min at RT. After stopping the reaction with IMDM media, cells were passed through a 40µm-filter, centrifuged at 300g for 6min and finally resuspend in PBS/1mM EDTA/1% FBS solution before counting. To identify the immune populations, 1.10^6^ cells were blocked in PBS/1mM EDTA/1% FBS/1µg of TrueStain FcX™ (Biolegend, 101320).

Thyroid lobes (8 lobes/condition/experiment) were cut in 1mm^3^ pieces and processed into single cells suspensions with the Mouse Tumor Dissociation Kit (Miltenyi Biotec, 130-096-730) diluted in Ca^2+^ free DMEM (Gibco, 21068-028) for 40 min at 37°C with up & downs with a P1000 every 8 min. Then, the cell suspension was passed through a 40 µm-filter and diluted in Ca^2+^ free DMEM /20% FBS/1mM EDTA pH8 to stop dissociation. Cells were centrifuged and resuspended in the same blocking solution as for splenic, thymic and blood cells for 10 min.

All cell samples were stained with anti-CD45, anti-CD11b, anti-CD3, anti-CD4, anti CD8a, anti-CD19 and anti-CD49b fluorochrome-conjugated antibodies. Dead cells were visualized by DAPI staining. Data were acquired on the BD LSRFortessa™ Cell Analyzer and analysed with FlowJo™ software (BD Biosciences). All antibodies and working conditions are recapitulated in Supplementary Table S1.

### 9. Western blotting

Western blotting was performed as described [28]. Briefly, lymph nodes from *Stabilin-1* WT (+/+) and *Stabilin-1* KO (-/-) mice were lysed in RIPA buffer and the protein concentration was measured using the bicinchoninic acid assay. 50µg of protein were loaded and electrophoresed through a 4-15% Mini-PROTEAN TGX precast gel (BIO-RAD, 4561084). Proteins were transferred onto a PVDF membrane, and the latter was blocked in TBS Tween 20 0.05% (TBST)/5% milk and incubated overnight with primary antibodies at 4°C (Supplementary Table 1). After washes in TBST, HRP-conjugated secondary antibodies were incubated in TBST/0.5% milk for 1 hour. Immuno-reactive bands were detected using Super Signal Chemiluminescent Substrate (Thermo Scientific) and images acquired using Fusion Solo S (Vilber Lourmat).

### 10. Statistical analysis

All graphs and statistical analyses were performed with Prism software (GraphPad Software) and are expressed as mean ± standard deviation. Real-time qPCR values were obtained by the ΔΔCT method [29]. Each graph represents the results obtained from a minimum of four different mice. Nonparametric statistical tests were used: Kruskal-Wallis followed by Dunn’s post-test for multiple comparison and Mann-Whitney for double comparison. Differences were considered statistically significant when p<0.05 (*); # stands for p<0.01; $ for p<0.001.

## RESULTS

### 1. Induction of BRAF^V600E^ expression in thyrocytes triggers a rapid increase in immune signalling pathways

Expression of the BRAF^V600E^ oncogene was obtained by intraperitoneal doxycycline injection in the Tg-rtTA/tetO-BRAF^V600E^ mouse model and led to tissue changes similar to those found in human PTC [22]. In our previous work, we observed (i) a two-fold increase in thyroid lobe size, (ii) the appearance of papillae in colloidal spaces, (iii) a progressive decrease in the protein reserve contained in the colloid, as well as (iv) a decrease in the expression of thyrocyte molecular markers, as from Day 1 to Day 4 (REF22). Additionally, a stromal reaction appeared from Day 2 onwards with progressive filling of the inter-follicular spaces with cells and development of a dense cellular sleeve around the thyroid lobe by Day 4 (Figure 1A).

**Figure 1:**
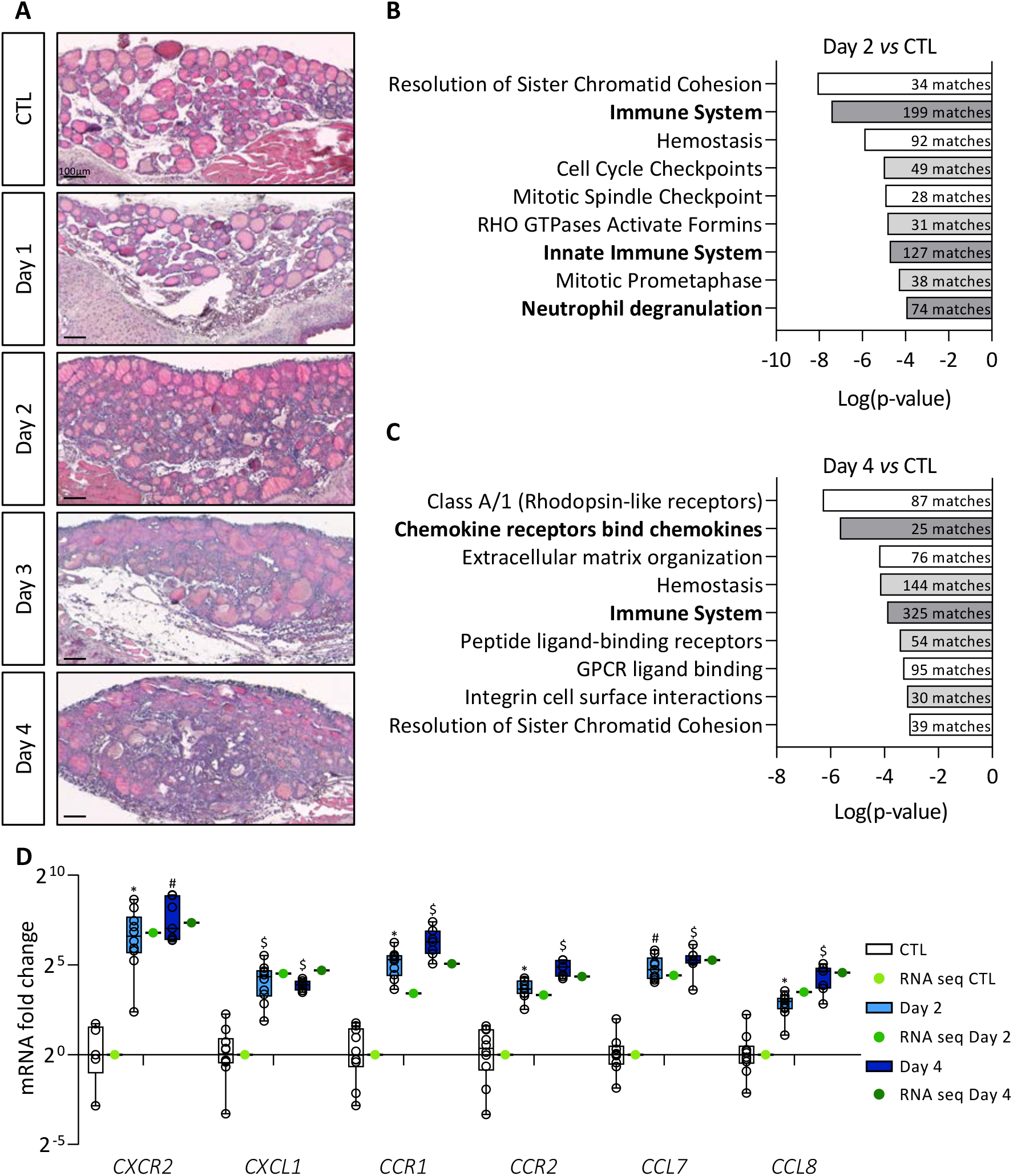
BRAF^V600E^ induction in mouse thyrocytes triggers stromal reaction which correlates with major changes in immune signaling pathways. (A) Hematoxylin-eosin staining of thyroid tissue from control (CTL) and doxycycline-treated mice, from Day 1 to Day 4. (B-C) Enriched signalling pathway revealed by MouseMine software using differentially expressed genes from total RNA sequencing (B) of Day 2 (n=4) compared to CTL (n=4) thyroids and (C) of Day 4 (n=4) compared to CTL (n=4) thyroids. (D) mRNA fold change of the most deregulated chemokines (*CXCL1, CCL7, CCL8*) and their receptors (*CXCR2, CCR1, CCR2*) found in RNA sequencing data (green dots) and measured by RT-qPCR (white and blue boxes) of CTL (n≥5) and doxycycline-treated mice, after 2 (n=11) and 4 days (n=7) of injections (Day 2, Day 4). mRNA fold changes are normalized on the geometric mean of *Rpl27* and *Gapdh* expression and compared to control group. Data are expressed as mean ± SD (* p<0.05, # p<0.01, $ p<0.001).

In order to understand the TME changes driven by BRAF^V600E^ mutation, we first performed a transcriptomic analysis. Sequencing of total RNA from control (CTL) and BRAF^V600E^ thyroids (Day 2 and Day 4) revealed more than 6000 differentially expressed genes (adjusted p-value <0.05) between Day 2 and CTL thyroids, and more than 9800 differentially expressed genes (adjusted p-value <0.05) between Day 4 and CTL (data not shown). We eliminated genes with log2 fold change between 2 and -2, and the remaining 1269 (Day 2 *vs* CTL) and 2716 (Day 4 *vs* CTL) genes were analyzed with MouseMine, a pathway enrichment analysis software (Figure 1B-C). After two days of BRAF^V600E^ induction, we observed enrichment of several cell cycle-as well as immune-related signaling pathways (Figure 1B). After four days of induction, we found enrichment of pathways associated with homeostasis and a persistence of those linked to the immune system (Figure 1C). Interestingly, immune-related signaling pathways showed the highest number of gene matches (numbers in dark bars of Figure 1C). Among the most deregulated genes at Day 4, we found many chemokines and their respective receptors. We confirmed the sequencing data by measuring the expression of some immune-related genes by RT-qPCR in tissues after two and four days of doxycycline injections (Figure 1D). Relative expression levels measured by RT-qPCR (blue boxes) were similar to those obtained by sequencing (green dots). Furthermore, we observed an increase in the expression of *CXCR2, CCR1*, and *CCR2* receptors, as well as of their ligands *CXCL1, CCL7*, and *CCL8* over the course of doxycycline treatment.

Taken together, these results revealed that induction of BRAF^V600E^ oncogene in thyrocytes caused a rapid stromal response in the thyroid and that a major aspect of this response is related to immune system signaling pathways.

### 2. BRAF^V600E^ induction triggers recruitment of macrophages, among which a population of CD11b+/LYVE1+/CD206+/STAB1+ immunosuppressive macrophages

Since stromal cellular density increased and since CXCR2, CCR1, CCR2 receptors are found on the surface of macrophages or myeloid-derived suppressor cells (MDSC), we studied macrophage recruitment in the thyroid upon BRAF^V600E^ induction. Immunostaining for F4/80 protein, a general macrophage marker, revealed a massive recruitment from Day 2 onwards (Figure 2A). At Day 4, each follicle appeared to be closely surrounded by abundant macrophages. Since we observed morphological changes in the thyroid, such as the progressive disappearance of the colloidal content, macrophage recruitment could serve in the cleaning of the tissue. However, lysosomes (LAMP1+, white) of the F4/80+ macrophages did not contain thyroglobulin (TG, red) at Day 2, and only few of these macrophages exhibited colocalization of LAMP-1 with TG marker, at Day 4 (Supplementary Figure S1A).

**Figure 2:**
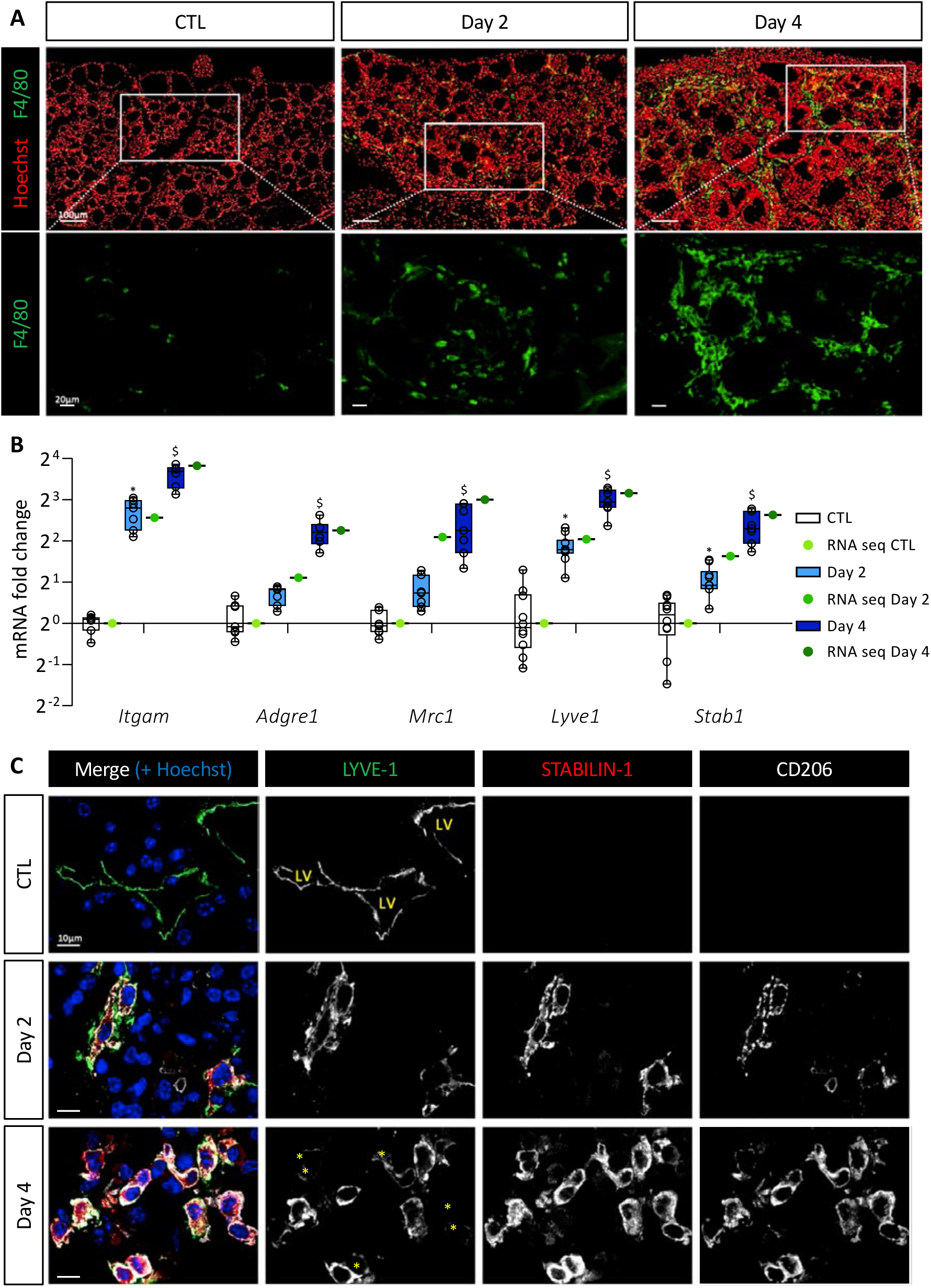
BRAF^V600E^ expression in mouse thyrocytes leads to massive macrophage recruitment including a particular macrophage population characterized by LYVE-1, CD206 and STABILIN-1 expression. (A) Immunohistofluorescence of F4/80 (green) on thyroid tissues from control (CTL) and doxycycline-treated mice (Day 2, Day 4). Hoechst (red) is used to label the nuclei. (B) mRNA fold change of different macrophage markers found in RNA sequencing data (green dots) and measured by RT-qPCR (white and blue boxes) from CTL (n=10) and doxycycline-treated mice, after 2 (n=11) and 4 days (n=7) of injections (Day 2, Day 4). mRNA fold changes are normalized on the geometric mean of *Rpl27* and *Gapdh* expression and compared to control group. Data are expressed as mean ± SD (* p<0.05, # p<0.01, $ p<0.001). (C) Immunohistofluorescence of LYVE-1 (green), STABILIN-1 (red) and CD206 (white) on thyroids from control (CTL) and doxycycline-treated mice (Day 2, Day 4). Hoechst (blue) is used to label the nuclei. LV indicates lumen of lymphatic vessels and yellow asterisks point cells with a very weak signal for LYVE-1 but positive for STABILIN-1 and CD206 markers.

We then analyzed by RT-qPCR and in the sequencing data the gene expression of myeloid cell markers, *i*.*e. Itgam* encoding for CD11b protein, and macrophage specific genes, i.e. *Adgre1* and *Mrc1* encoding respectively for F4/80 protein and CD206 protein, a marker of immunosuppressive macrophages. Expression of these different genes was increased after BRAF^V600E^ induction (Figure 2B). This increase was due to cell recruitment in the thyroid since the number of CD11b and CD206 immunolabelled cells increased from Day 2 (Supplementary Figure S1B). By RT-qPCR, we also noticed a quantitatively and dynamically similar increase in the expression of *Lyve1* and *Stab1* (Figure 2B). This was interesting since recruitment of F4/80+, CD11b+, LYVE-1+, STABILIN-1+ macrophages has been reported in a melanoma model [30]. Several publications have further demonstrated the presence of LYVE-1, a lymphatic vessel-specific glycoprotein, on macrophages in physiological [31] or pathological [30,32] conditions. STABILIN-1 was first described as a scavenger receptor found on the surface of lymphatic endothelial cells and venous sinusoids [33]. Subsequently, it was also described on the surface of some immunosuppressive macrophages under physiological [34] and pathological [35] conditions.

Co-immunolabeling of LYVE-1, STABILIN-1 and CD206 proteins in CTL thyroids revealed the presence of LYVE-1+ lymphatic structures, but the absence of STABILIN-1 and CD206 positivity (Figure 2C). However, after two days of BRAF^V600E^ induction, cells positive for the three markers were detected. The recruitment of those cells increased with time (Day 4) and further specification also occurred since, besides triple positive cells, some cells showed reduced or undetectable LYVE-1 marker (Figure 2C; marked with an asterisk in bottom panel).

Altogether, these results demonstrated a massive and rapid recruitment of macrophages following BRAF^V600E^ induction. A minority of these macrophages might be involved in endocytosis of proteins or cellular debris. On the contrary, a population of macrophages carrying immunosuppressive markers, such as CD206 and STABILIN-1 appeared in the thyroid two days after BRAF^V600E^ induction, and persisted in the tissue.

### 3. Generation and validation of a CRISPR-Cas9 Stabilin-1 knockout mouse

Several publications have demonstrated that STABILIN-1 expression on macrophages plays a role as an immunosuppressive protein [36]. In different models of subcutaneous allografts of cancer cells, tumors presented reduced size in the absence of STABLIN-1 [37,38], and decreased metastasis [37,39]. However, the role of STABILIN-1+ macrophages in the thyroid cancer or in an *in-situ* tumor model is unknown. We therefore took advantage of our *in situ* PTC model to study the role of these macrophages by combining CRISPR-Cas9 knockout of *Stabilin-1* with Tg-rtTA/tetO-BRAF^V600E^ transgenes.

We designed two guide RNAs to ablate the transcription initiation site as well as the entire first exon, containing the ATG start codon, out of the 69 exons of the *Stab1* gene (Figure 3A, scissors). These allowed removal of a DNA fragment of approximately 1300bp, as verified by PCR (Figure 3B). Amplicons generated with primers located outside and inside of the targeted regions (fw-rv-rv’), had a band size of ±2300bp and ±1300bp for the wild-type allele and ±1000bp for the deleted allele (Figure 3B, left). Amplification of the longest (±2300bp) product was highly variable (here, only visible in heterozygous (+/-) mice). A second forward primer (fwʹ), located within the first exon, was also used with the reverse primer (rv’) (Figure 3B, right). Amplification of a ±600bp band revealed the presence of at least one wild-type allele (in +/+ and +/-), while no amplification was possible in knockout mice (-/-) since both primers (fwʹ-rv’) were in the deleted region. *Stabilin-1* knockout embryos were obtained at the expected Mendelian ratio, and were indistinguishable from control littermates at adult age. Western blotting analyses revealed the presence of a 260kDa band in wild-type adult lymph nodes, which was undetectable in knockout organs (Figure 3C). Since we inactivated a gene expressed by a cellular population of the immune system, we analyzed the most important immune populations in blood, spleen and thymus of adult unchallenged mice by flow cytometry (Figure 3D, E, F). The percentage of the different immune populations was comparable in wild-type (*Stab1*^+/+^) and KO mice (*Stab1*^-/-^). The CRISPR-Cas9 *Stab1* inactivation thus led to a complete absence of STABILIN-1 full-length protein with no detectable changes in immune populations.

**Figure 3:**
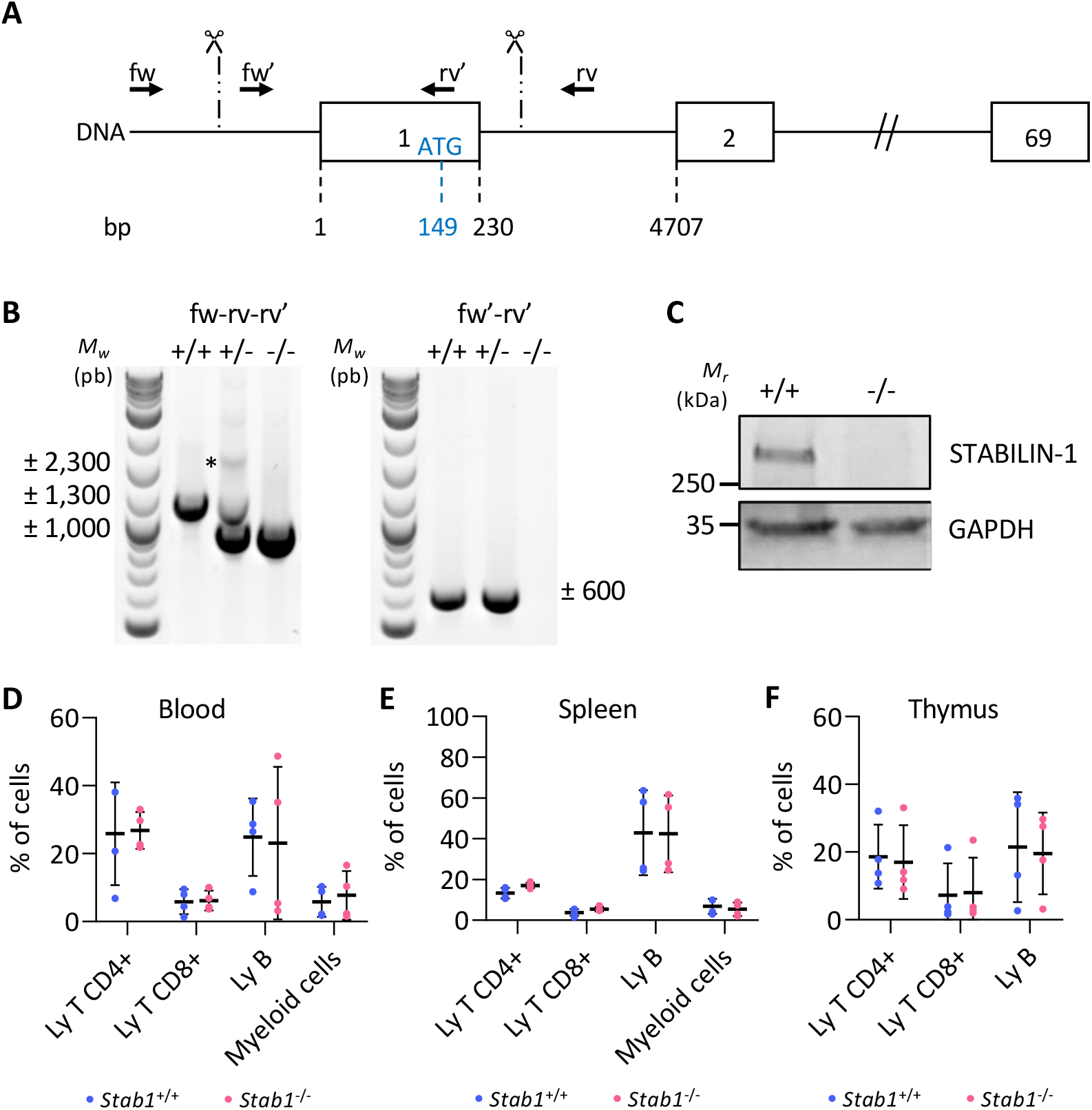
CRISPR-Cas9-based editing of *Stabilin-1* gene does not affect immune status of unchallenged mice. (A) Schematic representation of the *Stab1* locus with exons represented as boxes (only exons 1, 2 and 69 out of the 69 are illustrated). Numbers below indicate the position of the exon limits with respect to the transcription initiation site (+1). Position of the ATG (149 bases downstream of the +1) in the first exon is also indicated. Scissors and vertical dashed lines indicate the approximate region where the cuts occurred. Localization of genotyping primers are represented by black arrows. (B) Illustrative genotyping results for wild-type (+/+), heterozygous (+/-) and knockout (-/-) mice, obtained with two sets of primers. Asterisk points the absence of the theoretical wild-type band in +/+ mice. (C) Western blotting of STABILIN-1 protein in lymph node lysate from wild-type (+/+) and knockout (-/-) mice. GAPDH is used as a loading control. (D-F) Flow cytometry analyses of CD4+ T cells (CD45+, CD11b-, CD19-, CD3+, CD4+, CD8-); CD8+ T cells (CD45+, CD11b-, CD19-, CD3+, CD4-, CD8+); B cells (CD45+, CD11b-, CD19+); myeloid cells (CD45+, CD11b+) in blood (D) and from dissociated cells of spleen (E), and thymus (F) from wild-type (+/+) and knockout (-/-) mice (n=4).

### 4. Absence of STABILIN-1 does not affect epithelial PTC development in mice

To investigate if the absence of STABILIN-1 could impact on PTC development, wild-type (*Stab1*^+/+^) and KO (*Stab1*^-/-^) mice were treated with doxycycline every 24 hours for 4 days. Tissues were analyzed at Day 2 and Day 4. Hematoxylin-eosin staining (Figure 4A) revealed similar morphological changes following BRAF^V600E^ induction in *Stab1*^+/+^ and *Stab1*^-/-^ mice, *i*.*e*.: (i) increase in lobe size, (ii) progressive loss of colloid protein content, and (iii) appearance of papillae. Activity of the MAPK pathway, constitutively activated by the BRAF^V600E^ mutation, was assessed by measuring the expression of two target genes *Fosl1* and *Dusp5* by RT-qPCR (Figure 4B). Regardless of mouse genotype, expression of these two genes was strongly increased at Day 2, and slightly less at Day 4. Immunohistochemistry of phospho-ERK protein in the *Stab1*^+/+^ and *Stab1*^-/-^ thyroid lobes also confirmed the similar activation of the MAPK pathway as early as Day 2 and its maintenance at Day 4 (Supplementary Figure S2). Finally, we measured by RT-qPCR the expression of different genes involved in thyroid function, *Nis, Tpo, Tg, Tshr* as well as of two thyroid transcription factors, *Nkx2*.*1* and *Pax8* (Figure 4C). For *Nis, Tpo, Tg* and *Tshr*, the decrease was progressive. At Day 2 and at Day 4, expression of all these thyroid genes was decreased in a comparable manner in *Stab1*^+/+^ and *Stab1*^-/-^ mice. These results indicated that within four days the absence of STABILIN-1 did not impact on epithelial characteristics of BRAF^V600E^-induced PTC development.

**Figure 4:**
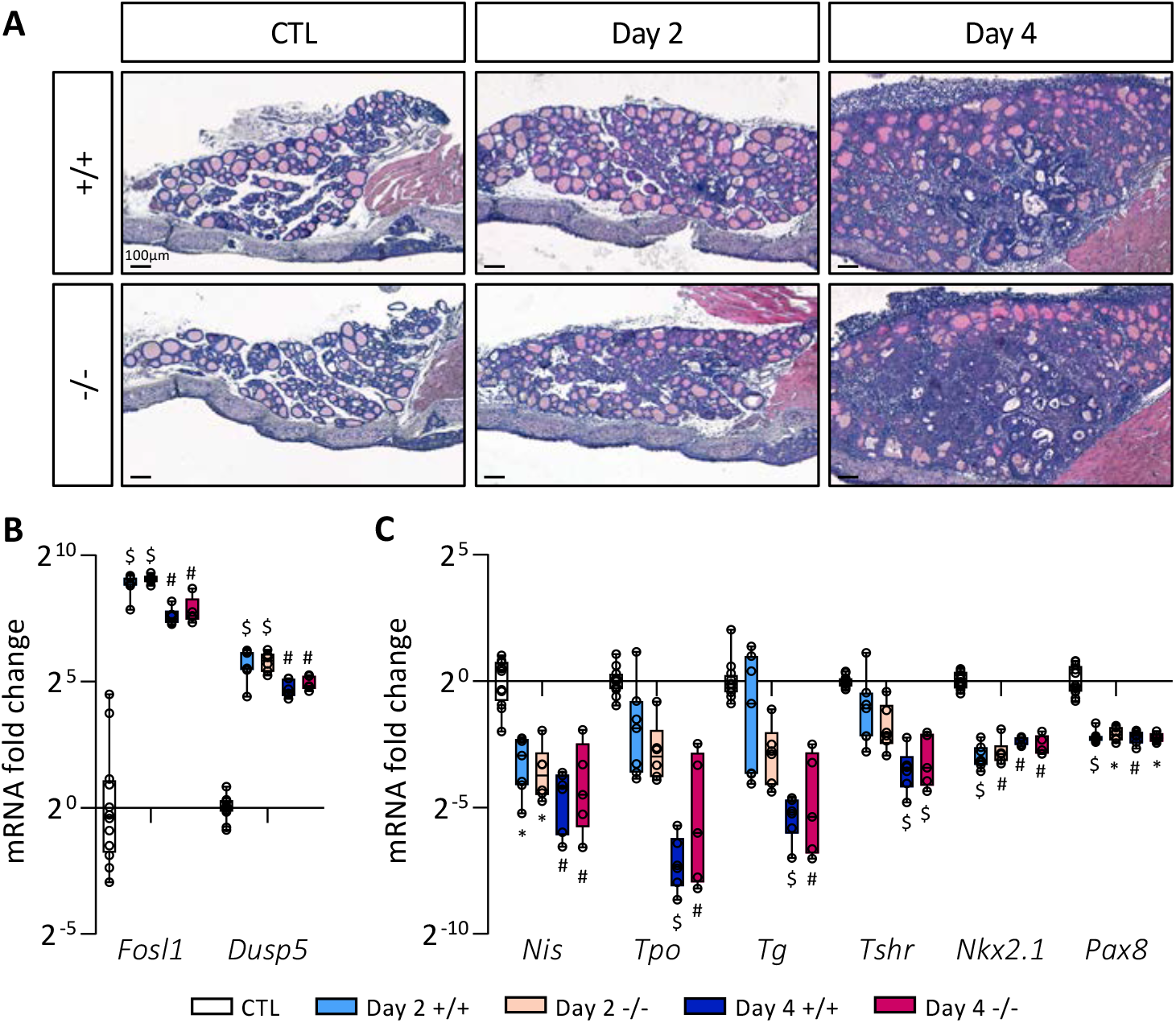
Absence of STABILIN-1 does not affect PTC development triggered by BRAFV600E expression. (A) Hematoxylin-eosin staining of thyroid tissue from wild-type (+/+) and knockout (-/-) mice treated with doxycycline (Day 2 and Day 4) or not (CTL). (B-C) RT-qPCR analyses of two MAPK pathway-target genes (b), and thyroid markers (C) in thyroid tissues from wild-type (+/+) and knockout (-/-) mice treated with doxycycline (Day 2 and Day 4) or not (CTL) (n≥5). mRNA fold changes are normalized on the geometric mean of *Rpl27* and *Gapdh* expression and compared to the CTL group (7 +/+ and 5 -/-thyroid samples). Data are expressed as mean ± SD (* p<0.05, # p<0.01, $ p<0.001).

### 5. Absence of STABILIN-1 does not affect circulating cell populations or macrophage recruitment in BRAF^V600E^-induced PTC

To assess circulating cell populations, we performed blood tests on *Stab1*^+/+^ and *Stab1*^-/-^ mice in CTL condition, at Day 2 and at Day 4 after BRAF^V600E^ induction. The numbers of white blood cells (WBC) and platelets were around 7×10^3^ and 700×10^3^ per mm^3^ of blood, respectively, in CTL (PBS-injected) *Stab1*^+/+^ mice. These numbers were not affected by BRAF^V600E^ induction at Day 2 and Day 4, nor by the absence of STABILIN-1 (*Stab1*^-/-^) (Supplementary Figure S3A). Similarly, the percentage of the different circulating cell populations was not affected by BRAF^V600E^ induction (Day 2 and Day 4 *vs* CTL), nor by the genotype (*Stab1*^-/-^ *vs Stab1*^+/+^), with ±75% of lymphocytes, ±3% of monocytes, ±12% of granulocytes, and ±7% of eosinophils (Supplementary Figure S3B). Finally, we measured and characterized, by flow cytometry, more specific circulating cell populations after 4 days of BRAF^V600E^ induction. After gating on live and CD45+ cells, we separated CD4+, CD8+ T cells, natural killer cells (CD19-CD49b+), B cells (CD4-, CD8-, CD19+) and myeloid cells (CD11b+) (Supplementary Figure S3C). Again, no change in abundance of these populations was observed in *Stab1*^-/-^ mice, as compared with *Stab1*^+/+^ mice.

We then studied macrophage recruitment in thyroid tissue 4 days after BRAF^V600E^ expression in *Stab1*^+/+^ and *Stab1*^-/-^ mice. We first measured by RT-qPCR the expression of the different macrophage markers analyzed previously, *Itgam, Adgre1, Mrc1* and *Lyve1* (Figure 5A). Expression of these genes was comparable in both *Stab1* genotypes. Conversely, using the same material, we observed in the *Stab1*^-/-^ mice a strong decrease (about 10-fold) of *Stab1* gene expression, not compensated by an increase in the expression of *Stabilin-2* gene (*Stab2*) (Figure 5A). Please note that despite a 10-fold decrease in *Stab1* gene expression, we did not observe a complete absence of transcript. This can be explained by the use of primers in exons 49 and 51 and an illegitimate transcription of the *Stab1* gene at low rate, despite deletion of the transcription initiation site in the knockout allele (Figure 3A). We then confirmed the presence of macrophages in the thyroid by immunolabeling (Figure 5B). In line with the RT-qPCR results, CD206+ cells were observed around the follicles in *Stab1*^+/+^ but also in *Stab1*^-^ /-mice, *i*.*e*. in the absence of STABILIN-1 staining.

**Figure 5:**
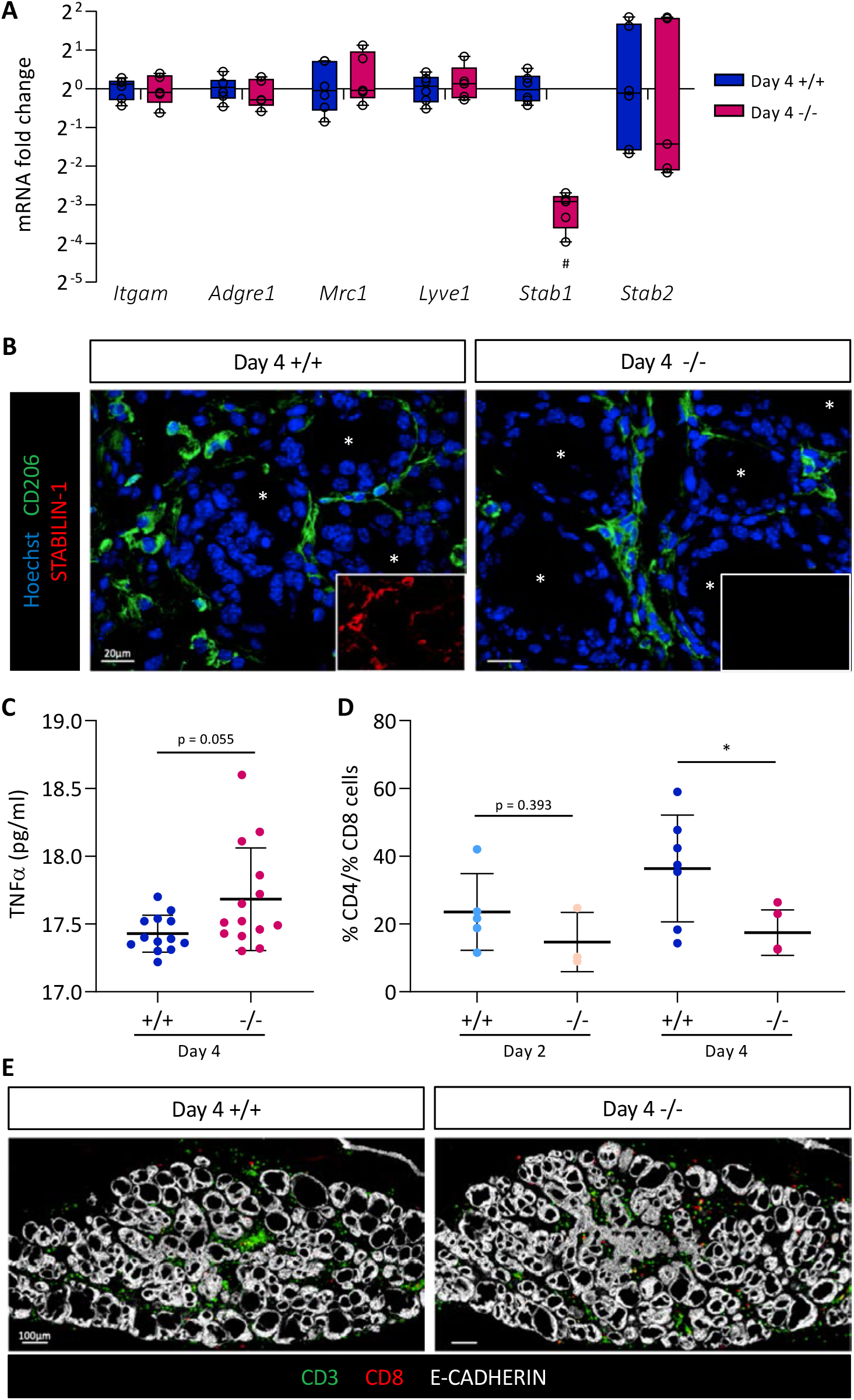
Absence of STABILIN-1 does not affect macrophage recruitment during PTC induction but changes the CD4+ T cells to CD8+ T cells ratio. (A) RT-qPCR analyses of macrophages markers and *Stabilin-2* (*Stab2*) genes in thyroid tissues from wild-type (+/+) and knockout (-/-) mice treated with doxycycline during 4 days (n≥5). mRNA fold changes are normalized on the geometric mean of *Rpl27* and *Gapdh* expression and compared to wild-type group (+/+). Data are expressed as mean ± SD (# p<0.01). (B) Immunohisto-fluorescence of CD206 (green) and STABILIN-1 (red, in small boxes) on thyroid tissue sections from wild-type (+/+) and knockout (-/-) mice treated with doxycycline for 4 days (Day 4). Hoechst (blue) is used to label the nuclei. White asterisks point the lumen of thyroid follicles. (C) Plasmatic TNFα concentration (pg/ml) measured by ELISA from wild-type (+/+) and knockout (-/-) mice treated with doxycycline for 4 days (Day 4). (D) Ratio of CD4+ T cells to CD8+ T cells calculated after the quantifications of immunohistofluorescently-labelled CD3 and CD8 cells in thyroid tissue sections from wild-type (+/+) and knockout (-/-) mice treated with doxycycline (Day 2 and Day 4). CD4+ T cells are considered as CD3+ CD8-cells. Each point represents the mean number of cells quantified from minimum three different (non-adjacent) sections of the same lobe (n≥4, * p<0.05). (E) Illustrative immunofluorescence, used for cell quantifications, of CD3 (green), CD8 (red) and the epithelial marker (E-CADHERIN, white) on thyroid tissue sections from wild-type (+/+) and knockout (-/-) mice treated with doxycycline for 4 days (Day 4).

Finally, we evaluated the expression profile of different markers and cytokines of pro-inflammatory (*TNFα, Nos2, Il1b* and *Il6*) and anti-inflammatory (*Mrc1, Arg* and *Il10*) macrophages in CTL, Day 2 and Day 4 thyroid tissue in *Stab1*^+/+^ and *Stab1*^-/-^ mice (Supplementary Figure S3D). We found a rapid increase in the expression of pro-inflammatory cytokines (*TNFα, Il1b* and *Il6*) two days after BRAF^V600E^ induction. Conversely, there was a slight delay in the response for anti-inflammatory macrophage markers and cytokines (*Mrc1, Arg* and *Il10*) which only displayed a significant increase at Day 4. These changes in gene expression were independent of the *Stab1* genotype (Supplementary Figure S3D). In a subcutaneous allograft model, Viitala *et al*. showed that the plasma concentration of TNFα was increased in *Stab1*^-/-^ mice [38]. We thus measured TNFα concentration in *Stab1*^+/+^ and *Stab1*^-/-^ mice, 4 days after *in situ* BRAF^V600E^-dependent PTC development. We also observed a trend towards increased plasma TNFα concentration in this *in situ* PTC mouse model, but without reaching significance.

Altogether, our data indicate that the absence of STABILIN-1 did not affect macrophage recruitment triggered by BRAF^V600E^ expression, but the increased trend in plasma TNFα concentration might suggest an increased pro-inflammatory response in the absence of STABILIN-1.

### 6. Absence of STABILIN-1 changes the CD4+/CD8+ T cell ratio in BRAF^V600E^-dependent PTC

It was demonstrated that blocking STABILIN-1 on macrophages or inactivating *Stab1* allowed the reactivation of CD8+ T cells within the tumor [38]. Thus, we first dissociated *Stab1*^+/+^ and *Stab1*^-/-^ thyroid tissue 4 days after BRAF^V600E^ induction, and quantified the percentages of CD4+ and CD8+ T cells from live/CD45+/CD19-/CD11b-/CD49b-DX5-cells (Supplementary Figure S4A) Out of 4 independent experiments, the ratio of CD4+ cells to CD8+ cells tended to decrease in *Stab1*^-/-^ mice compared to *Stab1*^*+/+*^ mice (Supplementary Figure S4B). However, this was not significant. Since these experiments required a lot of biological material (8 lobes/condition/experiment), we turned to immunohistofluorescence (Figure 5D-E). We used two lymphocyte markers, CD3 (green) and CD8 (red), in combination with epithelial cadherin (E-CADHERIN, white) to reveal the thyroid epithelium (Figure 5E). After nuclei segmentation, the number of each cell type was counted with the HALO software in CTL, Day 2 and Day 4 thyroid tissue in *Stab1*^+/+^ and *Stab1*^-/-^ mice. Due to the lack of a validated CD4 antibody working on sections, CD3+ CD8-cells were considered as CD4+ T cells. CD8+ T cells were CD3+ CD8+. CD4+ T cells were present in the thyroid before induction of BRAF^V600E^ expression and their number increased with time. CD8+ T cells were absent in CTL thyroids and were recruited during PTC development (Supplementary Figure S4C). Based on these quantifications, we calculated the CD4+/CD8+ ratio. At Day 2, the ratio of CD4+ T cells to CD8+ T cells was slightly decreased in *Stab1*^-/-^ mice (Figure 5D). This trend was confirmed at Day 4 when the ratio dropped from ±40 to ±20 in *Stab1*^-^ /-mice. This two-fold decrease was due to a slight decrease in the percentage of CD4+ cells and a slight increase in CD8+ cells in *Stab1*^-/-^ mice, as compared to *Stab1*^+/+^ mice (Supplementary Figure S4C).

Thus, the absence of STABILIN-1 caused a change in the lymphocytic populations present in the thyroid tissue, resulting in decreased CD4+/CD8+ T cell ratio, and supporting a potential immunosuppressive role for STABILIN-1 expression by macrophages recruited in the PTC thyroid tissue.

### 7. STABILIN-1+/CD206+ cells are present in benign and malignant human thyroid tissues

To extend our mouse data to patients diagnosed with thyroid diseases and in particular cancer, we investigated the presence of STABILIN-1+/CD206+ cells in benign tumors (multi nodular goiter, MNG, n=3), in healthy tissue neighboring a PTC (NHT, n=6) and in PTC (n=12) (Figure 6A). Interestingly, these STABILIN-1+/CD206+ macrophages were present and predominantly localized in the peripheral areas of the tumors for MNGs and PTCs. In MNG tissues, it seemed that the majority of STABILIN-1+ cells were not CD206+ while in PTCs we found different populations: STABILIN-1+ cells, CD206+ cells and CD206+ STABILIN-1+ cells (Figure 6A, tissue periphery). Within the tumor core, *i*.*e*. where E-CADHERIN density was very high, we found very few CD206+ and/or STABILIN-1+ cells (Figure 6A, tissue center). Lastly, in the neighboring healthy tissues (NHT), we found STABILIN-1+ cells with or without CD206 co-staining either within the epithelial thyroid tissue or at the periphery.

**Figure 6:**
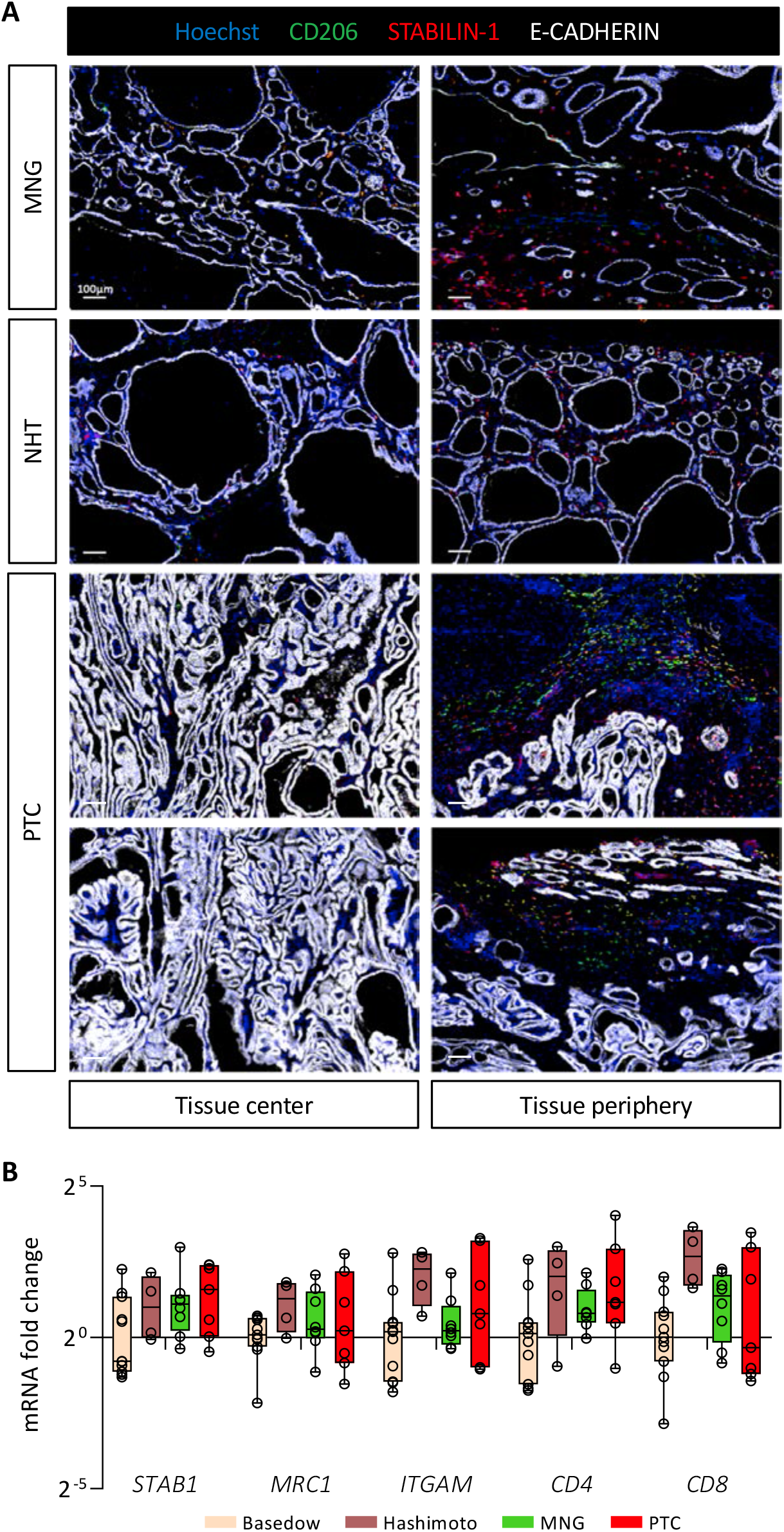
STABILIN-1+ cells are found in benign and malignant human thyroid tumors. (A) Immunohistofluorescence of CD206 (green), STABILIN-1 (red) and E-CADHERIN (white) on sections of human thyroid conditions: multinodular goiter (MNG), healthy tissue neighbouring of PTC (NHT) and papillary thyroid carcinoma (PTC). For each condition a picture in the center and the periphery of the tissue is shown. (B) RT-qPCR analyses of different immune cell population marker genes on different thyroid pathologies. mRNA fold changes are normalized on the *RPL13A* expression and compared to Basedow disease. Data are expressed as mean ± SD.

Finally, we measured by RT-qPCR the expression of different gene markers for immune cell populations in thyroid pathologies, such as autoimmune diseases (Basedow, n=11 and Haschimoto, n=4), benign (MNG, n=8) or malignant tumors (PTC, n=7) (Figure 6B). We found no statistical differences in the expression of genes related to immunosuppressive macrophages (*STAB1* and *MRC1*), myeloid cells (*ITGAM*) or T cells (*CD4* and *CD8*) in these pathological conditions. Of note, samples from Haschimoto’s patients seemed to display a higher expression for these different genes which correlates with its description as chronic lymphocytic thyroiditis [40]. This preliminary analysis revealed the presence of STABILIN-1+ cells in human diseased thyroid parenchyma, without showing a specific expression pattern.

## DISCUSSION

Although TC generally presents a 5-year overall survival rate higher than 95%, some tumors remain very aggressive and difficult to cure [6]. Among solid tumors with poor prognosis, it seems that beside the tumor cells, the TME and the cells which compose this environment also play a very important role in tumor progression [9]. By using a validated genetically-engineered, doxycycline-inducible mouse model of BRAF^V600E^-driven papillary thyroid cancer that mimics human PTC, we aimed to elucidate the changes in the tumor microenvironment induced by the BRAF^V600E^ mutation. Starting from a transcriptomic analysis, we identified major changes in immune signaling pathways associated with recruitment of a particular population of LYVE-1+/CD206+/STABILIN-1+ macrophages in the thyroid parenchyma. CRISPR-Cas9 inactivation of the *Stabilin-1* gene in the *in situ* PTC model revealed an increase in the number of CD8+ T cells in the thyroid TME during tumor progression. Finally, we also demonstrated the presence of CD206+/STABILIN-1+ cells in the parenchyma of benign and malignant human thyroid tumors.

The PTC model chosen for this study is a very powerful model. As soon as doxycycline is administered to mice, all thyroid cells start to express the BRAF oncogene. Moreover, the expression of the oncogene does not depend on its endogenous promoter but on a bacterial promoter, which causes strong and sustained transcriptional activation. Despite this non-physiological BRAF^V600E^ expression, this model has already made it possible to study MAPK pathway inhibitors that could be used as treatment [21]. Ryder *et al*. also highlighted the necessity of the CCR2 receptor for macrophage attraction and tumor development using this PTC model [19]. This PTC mouse model indeed presents several advantages: (i) ease of use as it only depends on the presence of two transgenes and the administration of doxycycline [41]; (ii) flexibility in the timing of oncogene induction and in the dose of doxycycline administered (either via food or via intra-peritoneal injections) [21,22]; (iii) short latency and high reproducibility as compared to sporadic thyroid cancer models [42-44]; and finally (iv), possibility to study early changes occurring in the thyroid which allows to investigate *in situ* PTC development as well as the thyroid immune microenvironment, impossible to study in xenograft models in immunosuppressed mice. Indeed, recruitment of TAMs described by Ryder *et al*. in human PTCs is correctly reproduced in this PTC mouse model [45]. Our RNA-seq data also support the comparative analysis of the TCGA and ESTIMATE databases performed by Zhao *et al*. which shows that the hub regulated genes are mainly related to immune signaling pathways with a majority of chemokine and cytokine genes identified [46].

The present study highlighted the recruitment of a particular population of immunosuppressive macrophages carrying LYVE-1/CD206/STABILIN-1 markers. This population has already been described in different types of tumor cell allografts from different organs (melanoma, lung carcinoma, lymphoma, colon cancer) [30,37,38]. In the melanoma allograft model, STABILIN-1 expression is induced on monocytes and on the tumor vasculature [37]. In the absence of STABILIN-1 on macrophages or vessels, tumor size is smaller and the number of metastases is lower. These changes are accompanied by a reduction in the number of CD206+ macrophages found within the tumor. In the lung carcinoma allograft model, tumors are also smaller in the absence of STABILIN-1 and this reduction in size is even more surprising if *Stabilin-1* is inactivated only on macrophages [38]. In this case, a decrease in the number of TAMs is observed but the TAMs present in the tumor express more MHC-II and CD206. In the PTC model used in this study, we did not observe STABILIN-1 expression on tumor vessels or changes in CD206+ macrophage populations upon *Stabilin-1* inactivation. This could result from the short read-out time (only 4 days). Indeed, absence of STABILIN-1 visualization on tumor vessels could be explained by the slower induction of STABILIN-1 expression on vessels as compared to TAMs, as described [38]. Alternatively, STABILIN-1 immunolocalization could be not sensitive enough to visualize small changes in protein expression on vessels. It would therefore be interesting to monitor *Stabilin-1* expression by *in situ* hybridization coupled with immunohistofluorescence to visualize the populations of interest [47,48].

The lack of tumor size reduction in the absence of STABILIN-1+ could also be explained by the short read-out of the model used. Indeed, we observed a tendency to decreased CD4+/CD8+ T cell ratio at day 2 and a significant decline after 4 days of doxycycline treatment. If the PTC mouse model could be analyzed at later time points, which is not, we could expect to observe a reduction in tumor size. However, we should not forget that STABILIN-1+ expression on lymphatic vessels has been shown to play an important role for lymphocyte trafficking to the draining lymph nodes and for dendritic cell entrance from the site of inflammation to the lymph nodes [49,50]. Thus, the total absence of STABILIN-1+ might impact on trafficking of these immune cells to and from the PTC, *e*.*g*. the CD8+ T cells which are mainly responsible for the decreased tumor size. We did not take the advantage of the macrophage-specific *Stabilin-1* knockout model (as in the study of Viitala *et al*. [38]) since it would have been complicated and time-consuming to implement in the *in situ* PTC model, already depending on the presence of two different transgenes Mechanistically, the role of STABILIN-1 macrophages in BRAF^V600E^-dependent *in situ* PTC is consistent with immunosuppression via their interaction with T cells and the change in the TNFα plasma concentration, as described in the different allograft models [38]. STABILIN-1 has also been proposed to be a scavenger receptor important for the clearance of extracellular tumor growth inhibiting molecules, such as SPARC or chitinase-like protein, from the TME in some tumor types (breast cancer, neuroblastoma), thereby leading to tumor growth [51-53]. Since these two STABILIN-1 ligands are rather pro-tumoral in human PTC [54,55], their clearance by STABILIN-1+ TAMs could therefore be detrimental for thyroid tumor growth.

In human, the expression level of *STABILIN-1* correlates with immunosuppressive environments such as the placenta or tumors [34]. In preeclampsia, STABILIN-1 expression on placental macrophages and circulating monocytes is decreased [56]. In different cancer (breast, head and neck, and colorectal), the presence of STABILIN-1 on tumor lymphatic vessels facilitates metastatic spreading to lymph nodes and its presence on intratumoral or peritumoral macrophages is associated with a poor prognosis [57,58]. The expression of STABILIN-1 on TAMs in urothelial bladder cancer is used as a biomarker since its presence is associated with increased mortality and poor response to chemotherapy [59,60]. The risk of recurrence of oral cavity cancer correlates with the density of STABILIN-1+ TAMs [61]. Finally, patients treated by immunotherapy with checkpoint inhibitors have a reduced response to the treatment if they express high level of *STABILIN-1* [36]. It is therefore important to consider *STABILIN-1* expression level in tumors to adjust the treatment. To the best of our knowledge, no study has described the presence of STABILIN-1+ cells in thyroid pathologies. Our preliminary analysis revealed its expression on TAMs in different pathologies but a larger number of samples as well as additional analyses are necessary to consider a role for STABILIN-1+ macrophages in thyroid pathologies. Future work should investigate the level of expression of *STABILIN-1* in different populations, such as vascular endothelial cells, lymphatic endothelial cells, and myeloid cells, present in the microenvironment of autoimmune or cancerous pathologies. It would also be relevant to analyze the correlation between *STABILIN-1* expression and the genetic status of the tumor such as the BRAF^V600E^ mutation, frequently found in PTCs and in melanomas where the presence of STABILIN-1+ TAMs has also been described [30].

## CONCLUSIONS

Our work has demonstrated rapid immune changes occurring in the thyroid parenchyma following expression of the BRAF^V600E^ oncogene in thyrocytes. This implicates the recruitment of a particular population of STABILIN-1+ immunosuppressive macrophages never described before in PTC. The absence of this protein changes the immune status of the tumor with the recruitment of CD8+ T cells into the tumor. It would thus seem that, as in allograft models, STABILIN-1+ TAMs play an immunosuppressive role in *in situ* PTC progression.

## Authors contributions

CS conceived and performed all the experiments with the help of PL, prepared the figures and wrote the manuscript. OD participated in data collection and analysis. ND and FH helped for flow cytometry experiments and analyses. AL analysed the mRNA sequencing. JK and JF developed and provided the Tg-rtTA; BRAF^V600E^ mice. YA generated the CRISPR-Cas9 *Stabilin-1* edited mice. MH and SJ shared the STABILIN-1 antibody and provided advices about STABILIN-1 data analysis. SL and PVM helped for the immune characterization of *Stabilin-1* knockout mice and for CD3-CD8 immunohistofluorescence. CD, MM and EM were involved in the human tissue sample collection. PH and DT advised and revised the manuscript. CEP and CS conceived and designed the study. CEP supervised the project and wrote the manuscript. All authors have approved the final version of the manuscript.

## Acknowledgements

The authors thank Romeo Delemazure and Abdelkadder El Kaddouri for technical help. CEP thanks Prof. Judith Varner and her group for advices and training.

## Institutional review board statement

The animal study protocol was approved by the the University Animal Welfare Committee, UCLouvain (2016/UCL/MD/005 and 2020/UCL/MD/011), Brussels, Belgium. Human tissue collection was conducted in accordance with the Declaration of Helsinki, and approved by the local ethics committee, UCLouvain (2017/25OCT/495 and 2017/04OCT/466), Brussels, Belgium.

## Informed consent statement

Informed consent was obtained from all subjects involved in the study.

---

**Supplementary figure 1:**
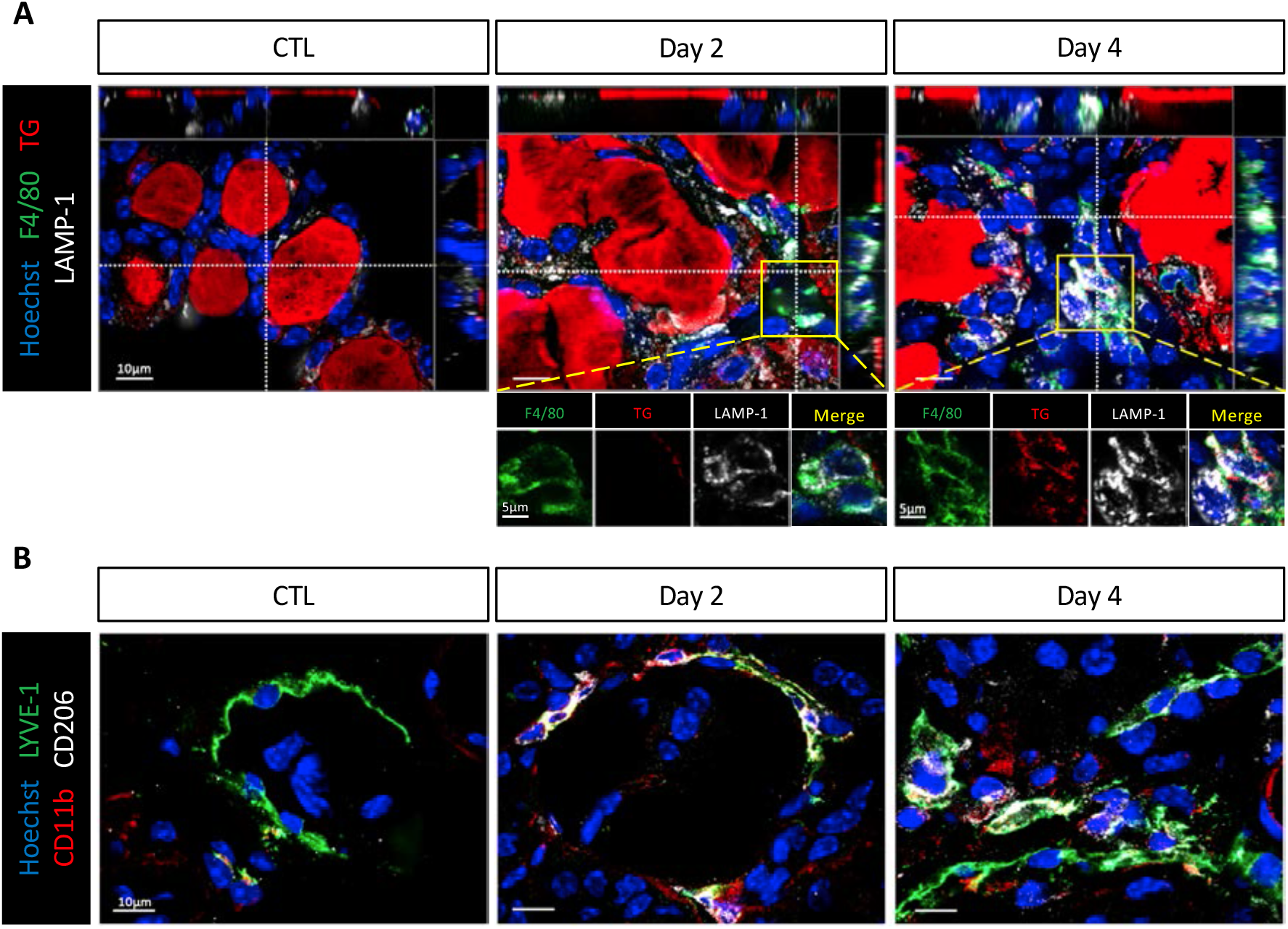
Macrophages are recruited following BRAF^V600E^ induction but do not display an important phagocytic activity. (A) Immunohistofluorescence of F4/80 (green), TG (red) and LAMP-1 (white) on thyroid sections from control (CTL) and doxycycline-treated mice (Day 2, Day 4). Hoechst (blue) is used to label the nuclei. Principal image is the result of a z-stack of 6µm (one picture per µm). Orthogonal lines refer to the selected focal section; x axis line is shown on the top and y axis line on the right of the principal image. Enlargements below of Day 2 come from another focal section than which shown on the principal image. (B) Immunohistofluorescence of LYVE-1 (green), CD11b (red) and CD206 (white) on thyroid control (CTL) and doxycycline-treated mice (Day 2, Day 4). Hoechst (blue) is used to label the nuclei.

**Supplementary figure 2:**
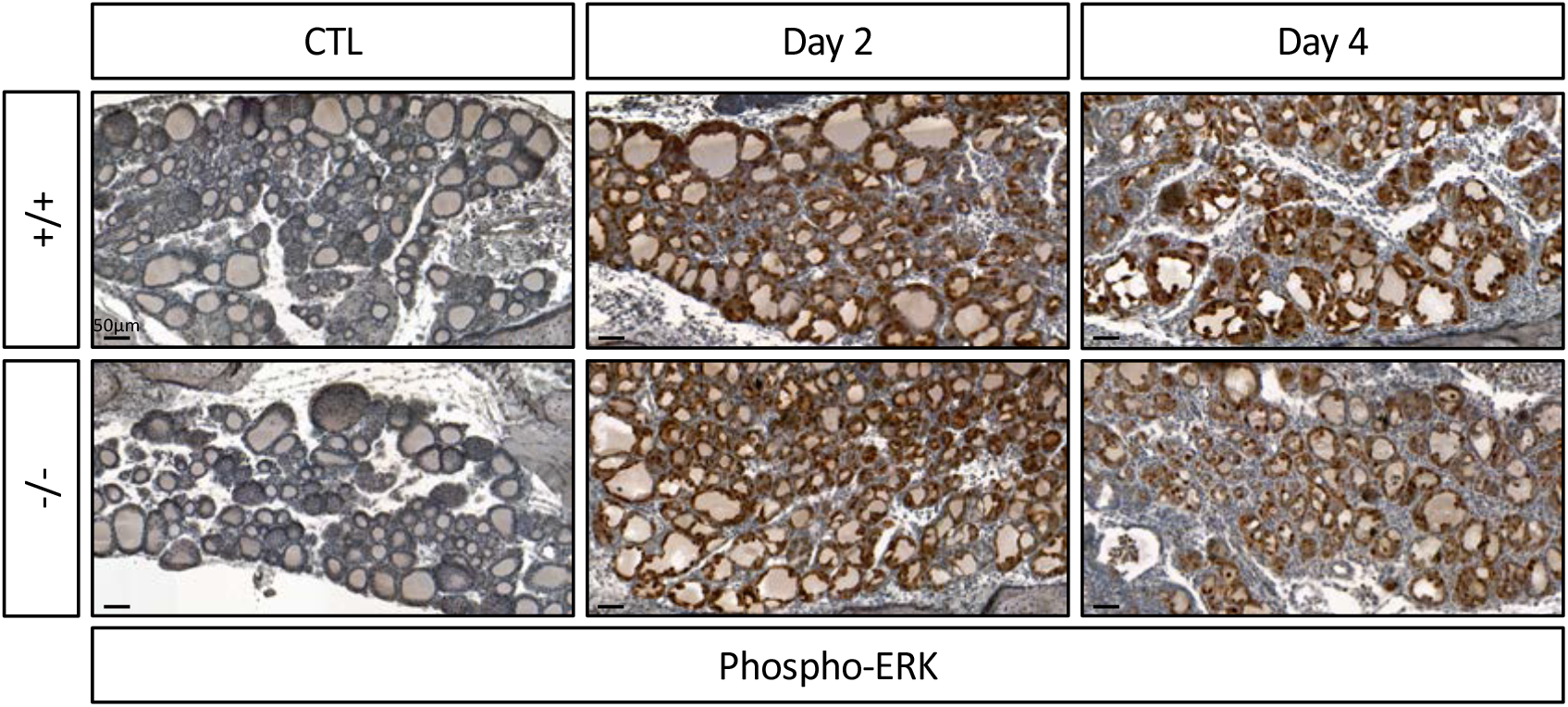
Absence of STABILIN-1 does not affect MAPK pathway activation in thyrocytes following BRAF^V600E^ induction. Immunohistochemistry of phospho-ERK on thyroid tissues from wild-type (+/+) and knockout (-/-) mice treated with doxycycline (Day 2 and Day 4) or not (CTL).

**Supplementary figure 3:**
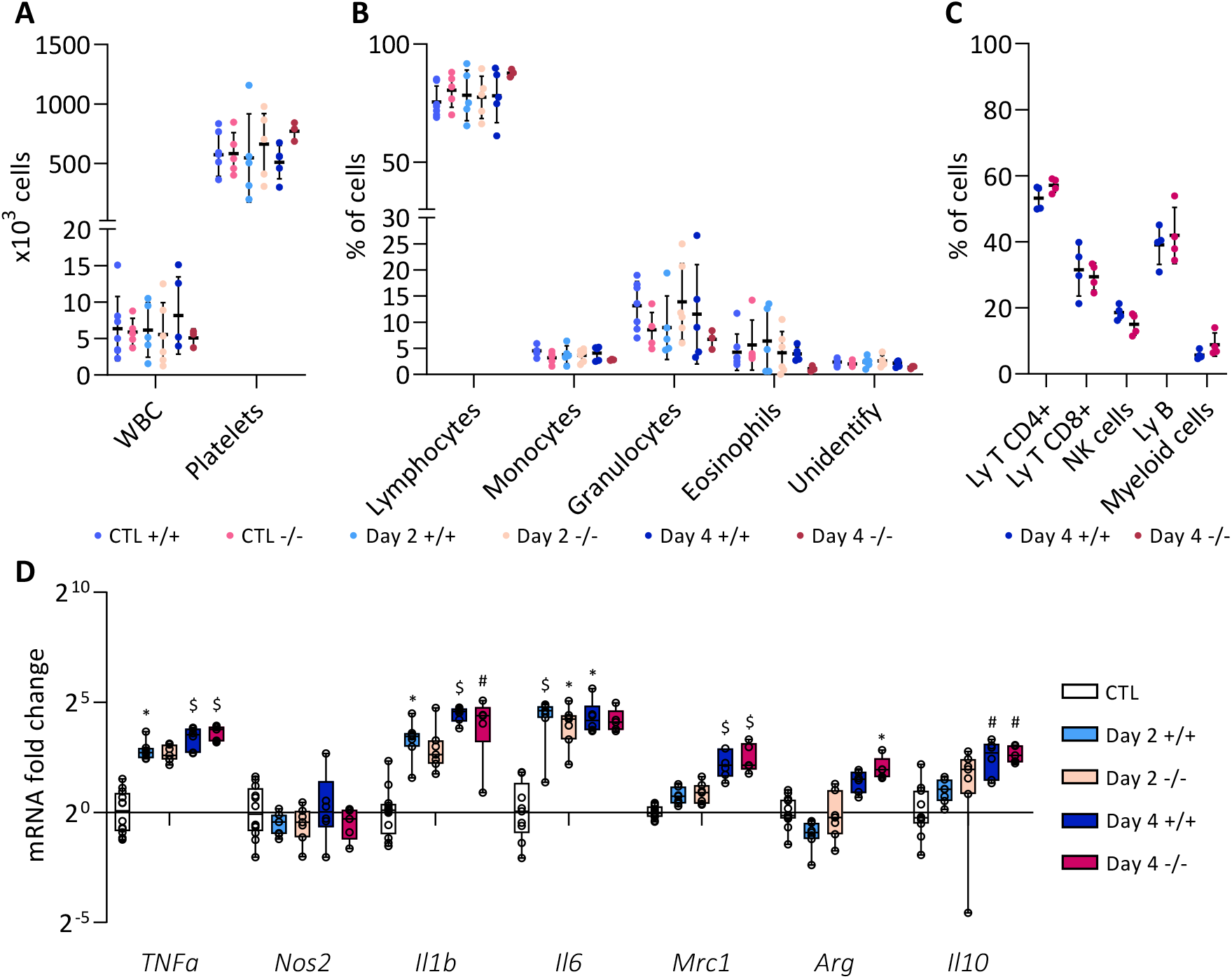
Absence of STABILIN-1 does not change the populations of circulating cells and does not impact macrophage markers. (A-B) Blood analyses of wild-type (+/+) and knockout (-/-) mice treated with doxycycline (Day 2 and Day 4) or not (CTL). (A) Total number of cells per ml of blood; (B) percentage distribution of white blood cells (n≥3). (C) Flow cytometry analyses of CD4+ T cells (CD45+, CD11b-, CD19-, CD49b-, CD3+, CD4+, CD8-); CD8+ T cells (CD45+, CD11b-, CD19-, CD49b-, CD3+, CD4-, CD8+); NK cells (CD45+, CD19-, CD49b+); B cells (CD45+, CD11b-, CD19+); myeloid cells (CD45+, CD11b+) in blood from wild-type (+/+) and knockout (-/-) mice treated with doxycycline for 4 days (Day 4)(n=4). (D) RT-qPCR analyses of pro-inflammatory macrophage markers genes (*TNFα, Nos2, Il1b, Il6*) and anti-inflammatory macrophage genes (*Mrc1*-already shown in Figure 5A-, *Arg, Il10*) from wild-type (+/+) and knockout (-/-) mice treated with doxycycline (Day 2 and Day 4) or not (CTL)(n≥4). mRNA fold changes are normalized on the geometric mean of *Rpl27* and *Gapdh* expression and compared to control group (7 +/+ and 5 -/-thyroid samples). Data are expressed as mean ± SD (* p<0.05, # p<0.01, $ p<0.001).

**Supplementary figure 4:**
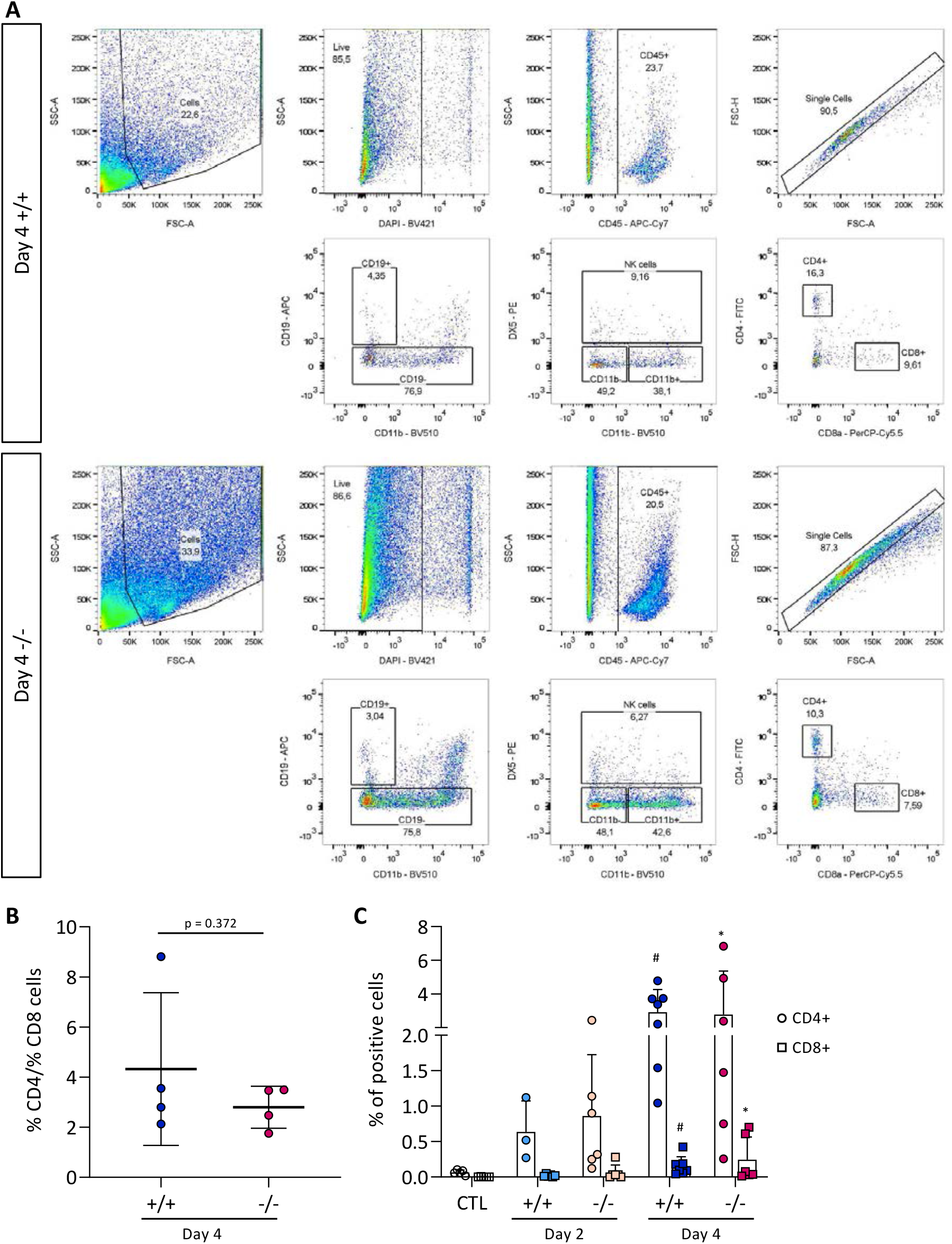
Absence of STABILIN-1 causes a slight increase of CD8+ T cells in thyroid parenchyma during PTC induction. (A) Illustration of the flow cytometry gating strategy to obtain CD4+ T cells to CD8+ T cells ratio of Supplementary figure 4B. (B) Ratio of CD4+ T cells to CD8+ T cells measured after flow cytometry analyses of dissociated thyroid tissues from wild-type (+/+) and knockout (-/-) mice treated with doxycycline for 4 days (Day 4). (C) Distribution (in percentage) of CD4+ T cells (round points) and CD8+ T cells (square points) of HALO quantifications on thyroid tissue sections from wild-type (+/+) and knockout (-/-) mice treated with doxycycline (Day 2 and Day 4) or not (CTL) (n≥5, compared to CTL condition; * p<0.05, # p<0.01).

**Supplementary table 1:**
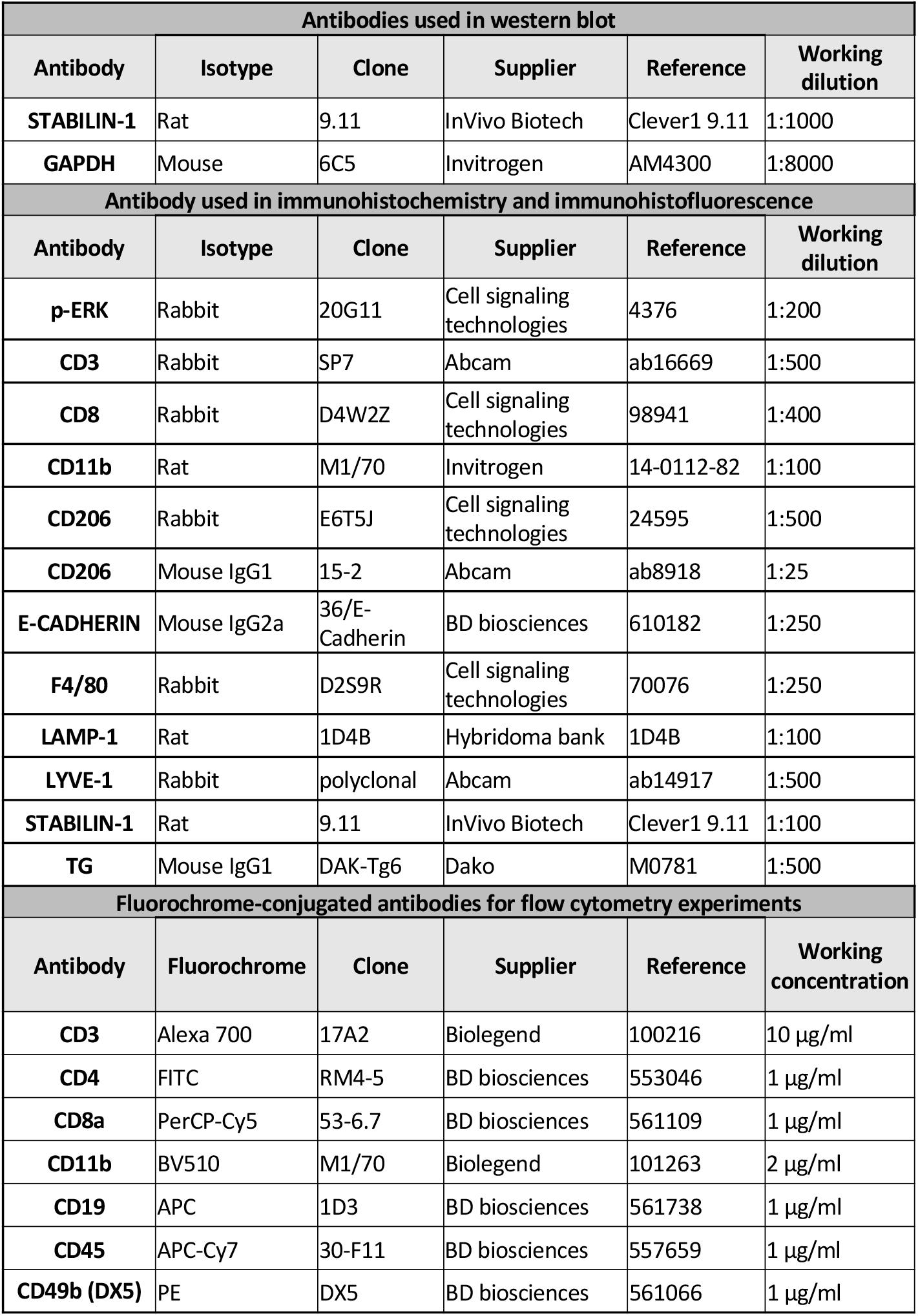
List of antibodies used

**Supplementary table 2:**
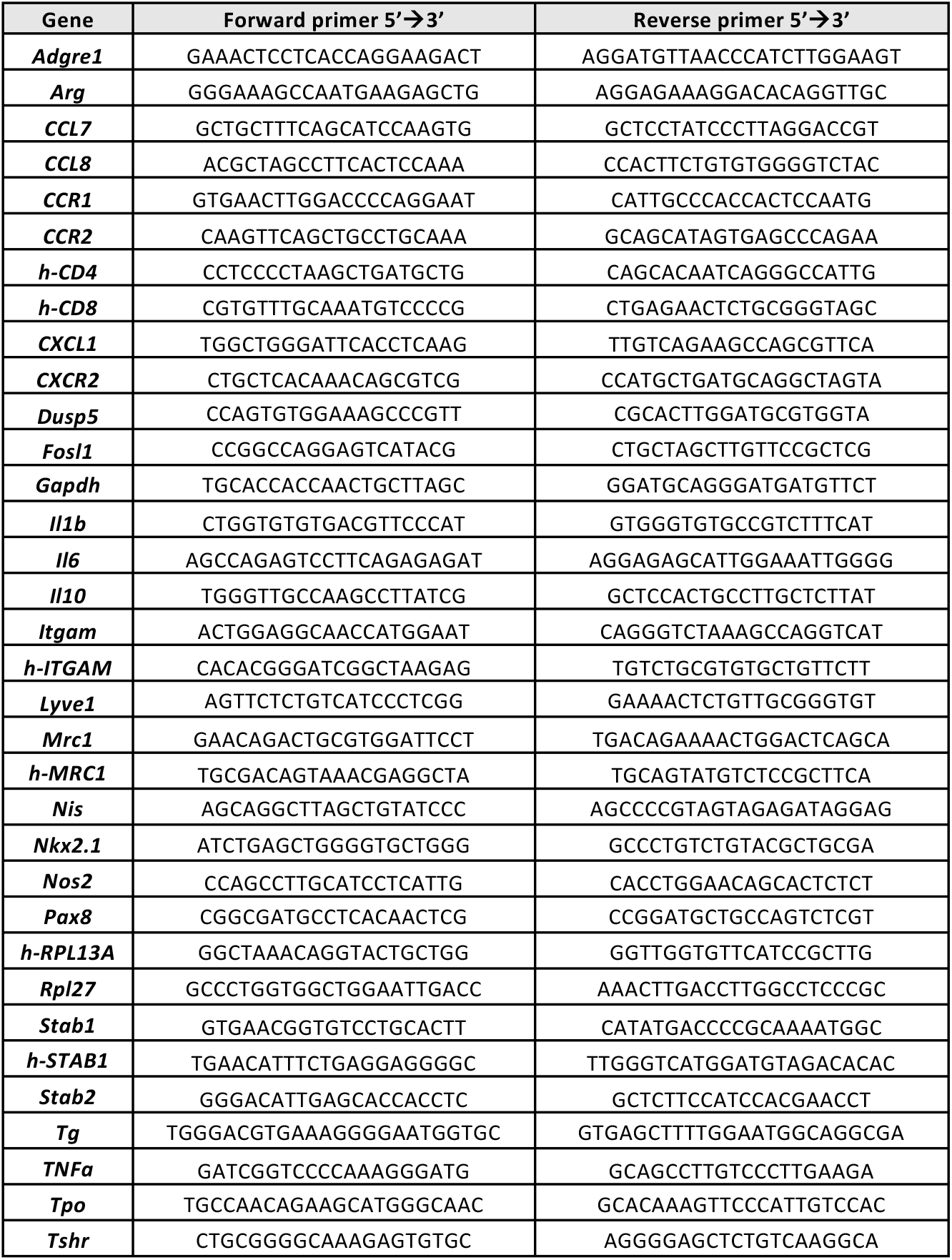
Sequences of primers used in mRNAs RT-qPCR

